# 3D test sample for the calibration and quality control of super-resolution and confocal microscopes

**DOI:** 10.1101/2020.10.14.336032

**Authors:** Ernest B. van der Wee, Jantina Fokkema, Chris L. Kennedy, Marc del Pozo, D.A. Matthijs de Winter, Peter N.A. Speets, Hans C. Gerritsen, Alfons van Blaaderen

## Abstract

A multitude of samples is required to monitor and optimize the quality and reliability of quantitative measurements of (super-resolution) light microscopes. Here, we present a single sample to calibrate microscopes, align their laser beams and measure their point spread function (PSF) in 3D. The sample is composed of a refractive index matched colloidal crystal of silica beads with fluorescent and gold cores. The microscope can be calibrated in three dimensions using the periodicity of the crystal; the alignment of the laser beams can be checked using the reflection of the gold cores; and the PSF can be measured at multiple positions and depths using the fluorescent cores. It is demonstrated how this sample can be used to visualize and improve the quality of confocal and super-resolution images. The sample is adjustable to meet the requirements of different NA objectives and microscopy techniques and additionally can be used to evaluate refractive index mismatches as a function of depth quantitatively.

---

Reliable quantitative 3D light microscopy measurements require a well calibrated setup. Moreover, proper alignment of the microscope contributes to a higher resolving power, thereby enabling analysis at a smaller scale^1^. Calibration and alignment have become even more critical with the advent of super-resolution microscopy techniques, such as stimulated emission depletion (STED) microscopy^2–4^. Additionally, it has been demonstrated that image restoration with an experimentally measured point spread function (PSF) allows for better image restoration than deconvolution with a theoretical PSF^1, 5–8^. Measurements of the PSF in 3D is also of importance if one wants to achieve the maximum possible accuracy in the analysis of microscopy images ^9, 10^. Therefore, it is important to be able to reliably measure the PSF in 3D of a microscope setup, in combination with a proper alignment and calibration in all three dimensions.

Currently, a multitude of different samples and methods are needed for a complete evaluation of a light microscopy setup that is to be used for quantitative measurements. First of all, calibration of the microscope in the lateral directions can be performed with commercially available stage micrometers. For calibration in the axial direction various methods have been reported^11–15^. In addition, the alignment of excitation beams is commonly checked with ∼100 nm gold beads^16^. For STED, the same gold beads can be used to check the alignment of the depletion beam with the excitation beam, which is crucial to maximize the performance of the system. Finally, measurements of the PSF of a confocal microscope can be performed by imaging sub-diffraction sized (< 175 nm in diameter) fluorescent beads^17^ or larger beads in combination with deconvolution software^14^ in 3D. For STED this is more challenging since smaller fluorescent probes (< 50 nm in diameter) are required, as the size of the probes needs to be close to, or below, the resolving power of the technique. Some suitable probes are available: DNA origami techniques for example have been demonstrated to produce sub-30-nm fluorescent probes for PSF measurements and the calibration of a STED microscope^18, 19^. In addition, it has been demonstrated that photostable quantum dots can be imaged with STED^20, 21^ and could therefore be used as fluorescent probes for PSF measurements of a STED microscope. However, a simple method to ensure enough separation between the probes in 2D or a method to produce 3D samples, with easily tunable refractive index contrast, is not yet available.

While the PSF measurement and alignment methods used so far are good tools to check the performance of a microscope, these methods usually only apply to measurements close to the cover glass. In recent work using (3D) STED in combination with glycerol objectives, super-resolution imaging far from the cover glass was demonstrated in life science specimens, where the refractive index is close to 1.45^1, 22, 23^. In addition, knowledge of the depth dependent PSF has been shown to be important for accurate 3D image reconstruction^8^ and high accuracy image analysis ^9, 10^.

In this work, a single sample for the 3D calibration and alignment, as well as the PSF measurement of a confocal (STED) microscope is presented. The sample is composed of a colloidal crystal consisting of a mixture of highly monodisperse silica beads with a gold or fluorescently labeled silica core. The crystal is refractive index matched with an embedding solvent mixture, resulting in optimal imaging conditions and an effective refractive index similar to life science specimens optimized for glycerol objectives. The periodicity of the crystal of touching particles in the lateral and axial directions is used as a ruler to perform calibrations, and is also geometrically stable against variations of the liquid that is in between the particles. The gold cores are used to image the excitation and depletion spots in reflection and to align the STED microscope. The fluorescent cores are individually resolvable as they are separated by a non-fluorescent silica shell and can therefore be used for PSF measurements at different positions and depths in the sample. By ensuring that the particles with the different cores have the same total size and the same inter-particle interactions, the resulting colloidal crystals are solid solutions of the different types of core-shell particles on the close-packed lattices.

## Results

### The sample

The sample used for calibration, alignment and PSF measurement is depicted in Fig. 1. The sample is composed of silica beads with an average diameter of 505 nm (< 2% polydispersity index (PDI), where PDI is the standard deviation over the average diameter) that are assembled into a colloidal crystal on a cover glass by vertical deposition^24^ (Fig. 1a). The thickness of the crystal can be tuned by performing repeated deposition steps or the concentration of particles during deposition. The cores of the silica beads consist of either gold (80 nm in diameter) or rhodamine B fluorescently labeled silica (45 nm in diameter). The number ratio between the two types of particles was chosen to be approximately 1 to 100, respectively.

**Figure 1:**
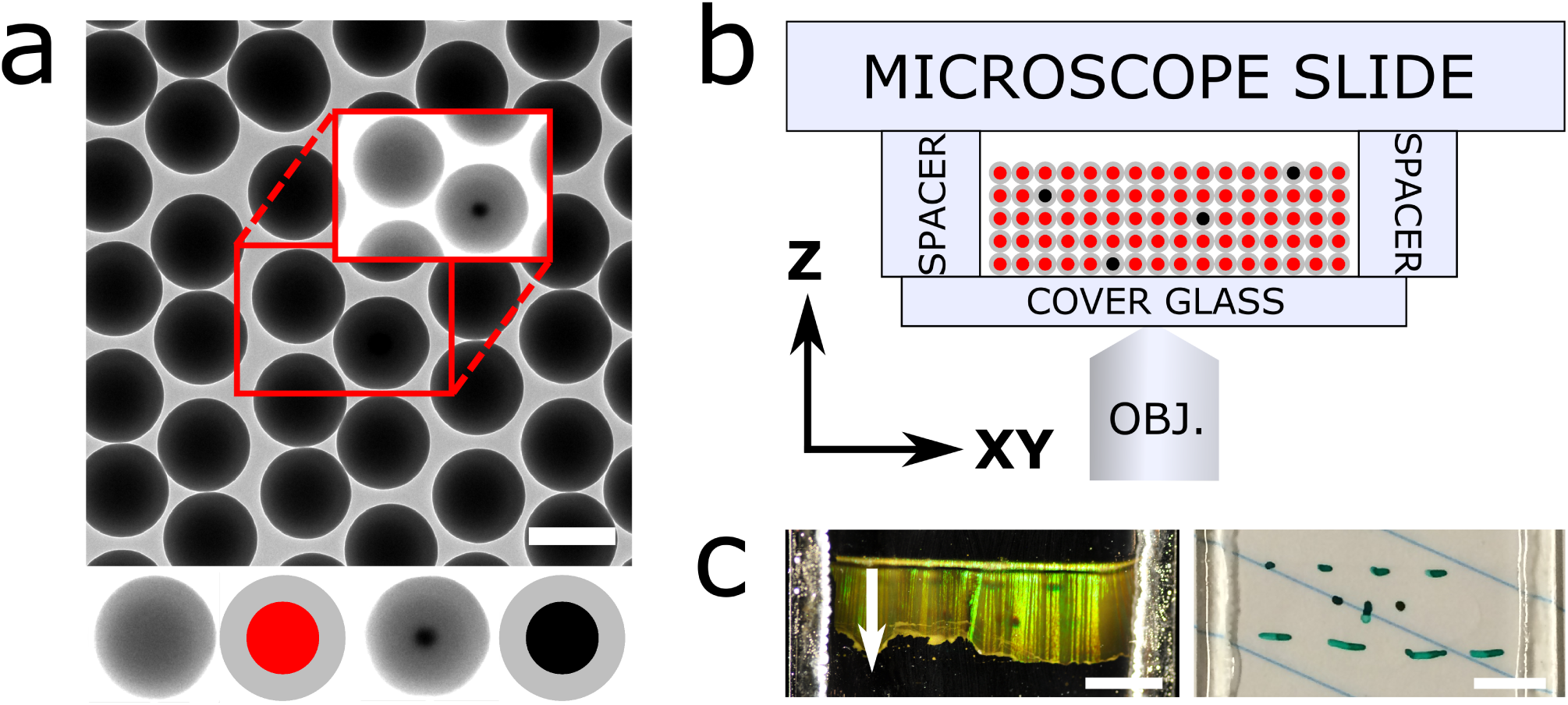
Calibration and alignment sample. a) Transmission electron micrograph (TEM) of the two building blocks of the sample: monodisperse silica-coated 45 nm fluorescent silica (red) and 80 nm gold (black) cores. The cores of the schematic particles are not drawn to scale. Inset: upon (TEM) contrast variation the gold cores become visible. The scale bar is 500 nm. b) Side view of the sample consisting of a crystal composed of the particles in *a)*, refractive index matched for optimal imaging. c) Top view of the sample showing the bright Bragg reflections of the index matched crystal upon illumination from the side (left) and full transparency under available light (right). The direction of crystal growth during vertical deposition is parallel to the *Y* direction. The scale bars are 0.5 cm.

To characterize the structure of the sample, a crystal grown using the same method, but with slightly larger particles was imaged using focused ion beam scanning electron microscopy (FIB-SEM) tomography^25, 26^ (see Supplementary Note 1). From the recorded FIB-SEM data the coordinates of the particles were determined, as described in ref. 26. From the coordinates we could identify the crystal structure as face-centered cubic (FCC) with stacking faults of hexagonal close-packed (HCP) order using bond orientational order parameters^27^. It was also observed that the hexagonal arrangement of the particles in the (111) plane of the FCC crystal is not perfectly hexagonal. In the direction of crystal growth (*Y* in Figure 1c) the distance to the nearest neighbor was ∼4% smaller than in the two other nearest neighbor directions of the hexagon. This is in agreement with earlier studies using X-ray diffraction and FIB-SEM tomography where a similar crystal growth method was used^26, 28^. This slightly anisotropic deformation of the close-packed layers is almost certainly caused by the strong capillary forces between the drying liquid and the particles and the orientation of the layers with respect to the drying front. In addition, the layer spacing of the crystal perpendicular to the (111) plane was constant throughout the crystal (see Supplementary Fig. 1).

The sample was prepared from the crystal as follows. A cover glass with the crystal was glued onto two glass spacers fixed onto a microscope slide (Fig. 1b). To suppress scattering during imaging of the sample and distortions of the PSF, the crystal was infiltrated with a refractive index matching solvent mixture composed of glycerol and *n*-butanol (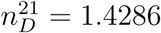, Supplementary Note 2). After infiltration, the sample was fully transparent under available light, whereas bright Bragg reflections were observed upon illumination from the side (Fig. 1c). The sample was sealed using UV glue to ensure a shelf life of at least, by the time of writing, several months.

### Lateral and axial calibration

A z-stack of the silica beads with rhodamine B labeled cores was recorded in fluorescent mode using a Leica HC PL APO 93×/1.30 GLYC motCORR STED WHITE objective (Fig. 2a). The periodicity of the crystal in both the lateral as well as the axial direction becomes visible after deconvolution of the z-stack (Fig. 2b-d). The (111) plane of the FCC crystal is parallel to the lateral plane, while the 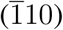 and 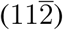 planes are parallel to the *XZ* and *YZ* directions, respectively, of the confocal z-stack. The growth direction of the crystal was parallel to the *Y* scanning direction of the confocal microscope.

**Figure 2:**
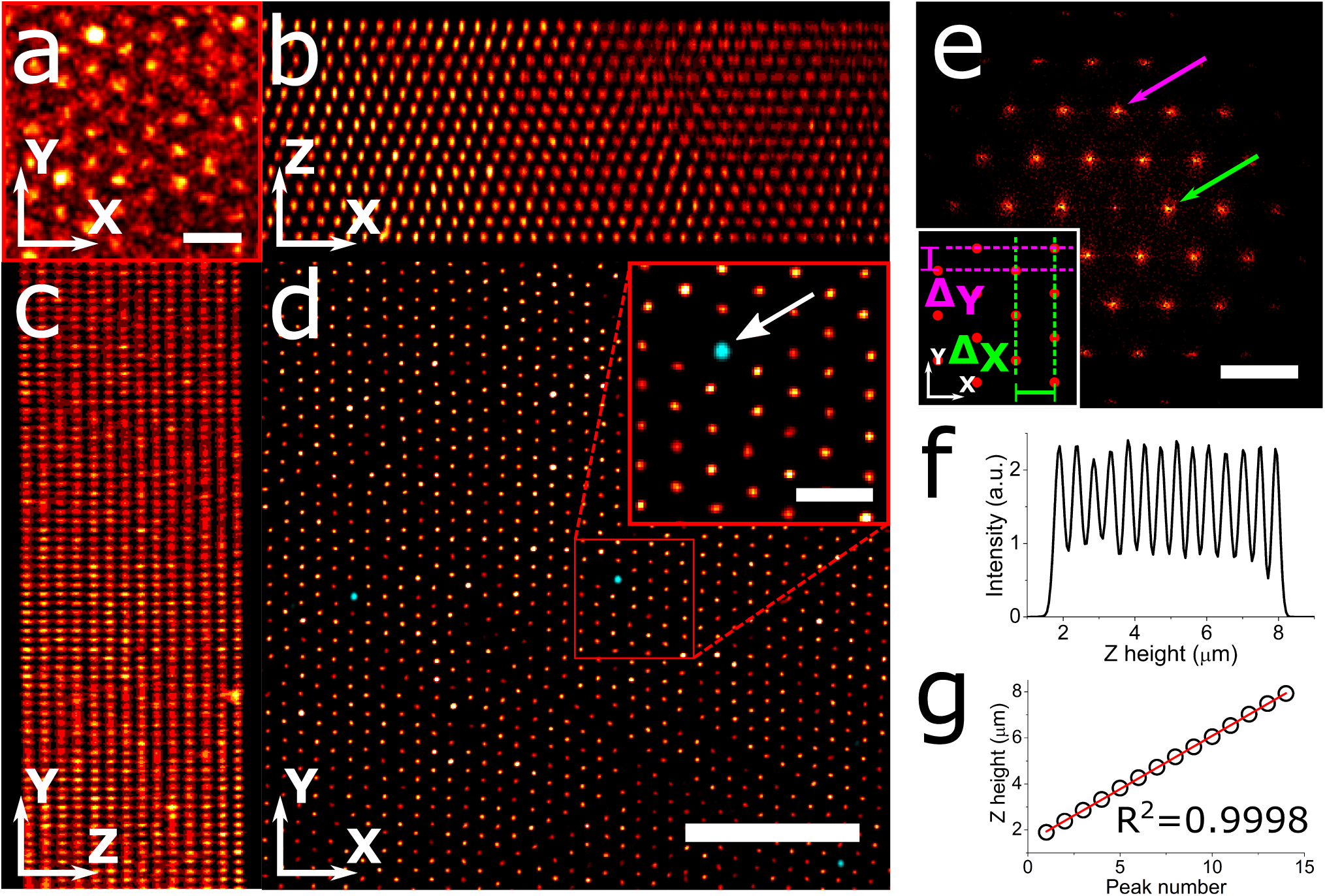
Calibration of confocal microscope. a) Close-up of a slice 3.6 µm from the cover glass from a confocal stack of the colloidal crystal (1 pixel Gaussian blur) acquired using a Leica HC PL APO 93 ×/1.30 GLYC motCORR STED WHITE objective and a pinhole of 0.7 Airy units. The scale bar is 1 µm. b) Average projection of the *XZ* planes of the confocal z-stack shown in (a) after deconvolution, showing the ABC stacking in the 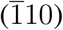 plane of the FCC crystal. c) Average projection of the *XY* planes of the deconvolved confocal z-stack, showing the 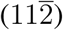 plane of the FCC crystal. d) Overlay of an *XY* slice of the confocal stack of both reflection (cyan) and deconvolved fluorescence (red) 3 µm from the cover glass. The arrow points at a gold core bead imaged in reflection mode. The scale bars are 5 and 1 (inset) µm. e) Fast Fourier transform of d) demonstrating long range hexagonal order. The arrows point to the peaks in the FFT image used for calibrating distances in the X (green) and Y (magenta) direction. The scale bar is 3 µm^−1^. The inset shows the characteristic distances Δ_*X*_ (green) and Δ_*Y*_ (magenta) in the realspace crystal corresponding to the peaks in the FFT image. f) Intensity profile in the z-direction, demonstrating the periodicity in the axial direction. g) Linear fit of the peak positions in f).

For calibration in the lateral direction the Fast Fourier transforms (FFT) were calculated of three *XY* slices from the deconvolved 3D confocal stack, respectively 1.3, 3.6 and 5.9 µm away from the cover glass. Figure 2e shows the FFT of the *XY* slice 3.6 µm away from the cover glass, with the arrows pointing at two characteristic distances in the (111) crystal plane in both the *X* (green) and *Y* (magenta) directions (see inset for the corresponding distances in real-space). From the three slices the characteristic distances Δ_*X*_ and Δ_*Y*_ were determined to be 481±4 nm and 269±2 nm, respectively. These values can be used to calibrate the two lateral directions of the microscope. Since the hexagonal arrangement is slightly distorted in the direction of crystal growth, *Y* in this case, the orientation of the hexagonal arrangement (and therefore the crystal growth direction) with respect to the confocal scanning direction should be taken into account during the calibration. As the particles are touching in the *Y* direction, the effective particle diameter *σ*_*eff*_ can be estimated by 2Δ_*Y*_. We find that *σ*_*eff*_ = 538±4 nm and therefore *σ*_*eff*_ ≈ 1.07*σ*_*TEM*_. This slightly larger effective diameter compared to the diameter as measured in TEM is expected because the particles shrink due to electron beam exposure during the TEM measurements^29^.

For the axial calibration of the setup, the intensity of the deconvolved confocal z-stack is plotted as a function of the *Z* height (Fig. 2f). The relative positions of the peaks in this intensity profile display almost perfect linearity (*R*^2^ = 0.9998, Fig. 2g) and a linear fit results in a distance of 448±2 nm between the lateral layers of the crystal. It should be noted that axial distances, as recorded by confocal microscopy using high NA objectives, are sensitive to refractive index mismatches between the sample medium and the immersion medium of the objective. This results in an apparent elongation or shrinkage of measured distances^14, 30^. The presented sample has a slightly lower refractive index (1.43) than the refractive index of the immersion liquid (1.45). Therefore, the axial scaling factor was determined using the method described by Besseling *et al*.^14^. A scaling factor of 0.982±0.003 was determined, resulting in a (*111*) layer spacing *d*_111_ of the sample of 440±2 nm. For a perfect FCC crystal *d*_111_ can be calculated using: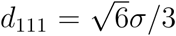, where *σ* is the particle diameter^28^. Using *σ*_*eff*_ = 538±4 nm, as estimated before, a value of *d*_111_ of 439±3 nm is found, which corresponds well to the measured value (*d*_111_ = 440±2 nm).

### PSF measurement and laser beams alignment

Using the fluorescent cores of the beads in the colloidal crystal, the PSF of a confocal microscope can be determined. Figure 3a shows the sub-resolution fluorescent cores close to the cover slip, imaged in confocal mode with a Leica HC PL APO 100×/1.40 OIL STED WHITE objective and the pinhole set to 0.7 Airy units. The PSF of this microscope has an expected ellipsoidal shape in the axial planes, but is slightly tilted in the *YZ* plane.

**Figure 3:**
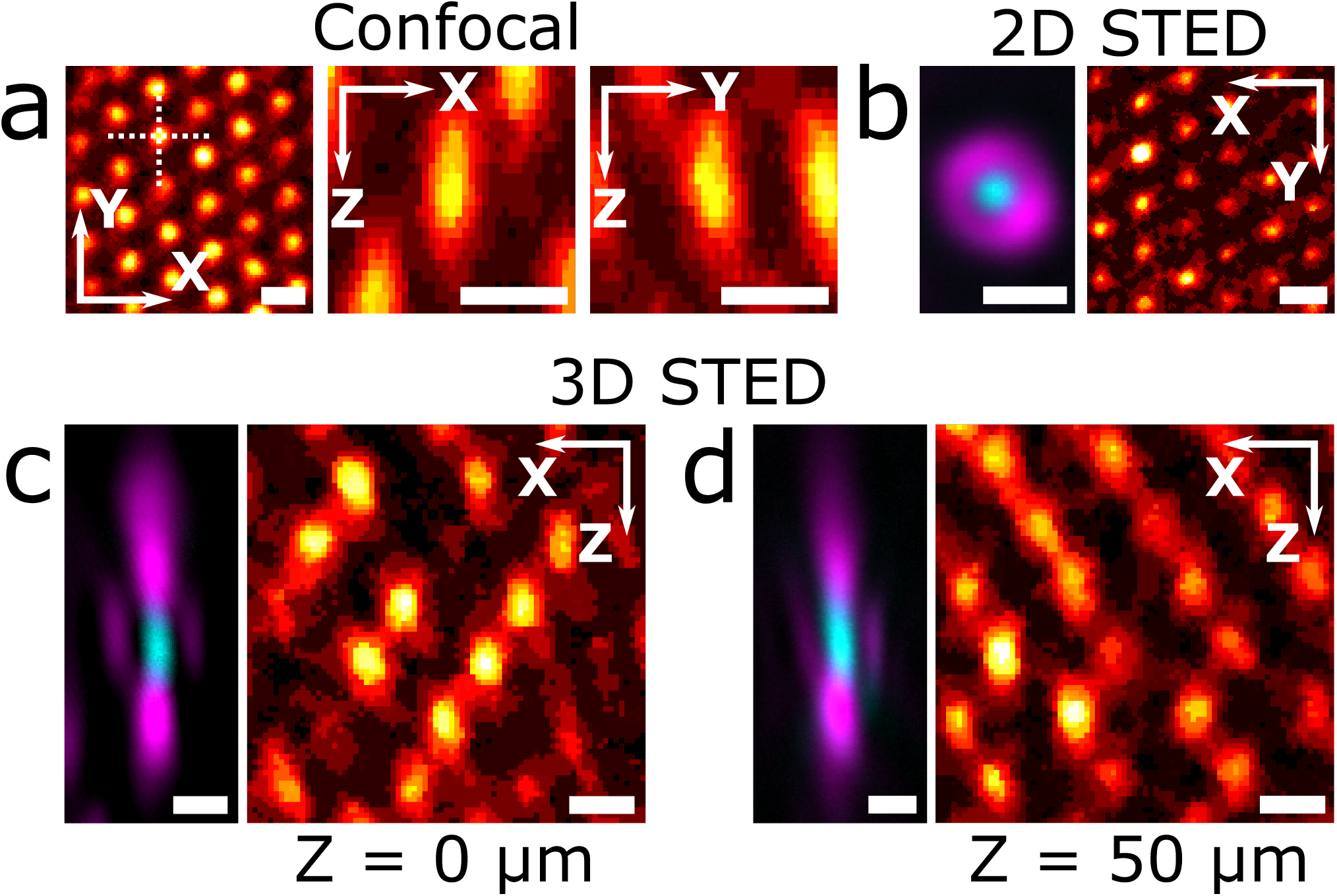
Alignment of confocal STED microscope. a) Slices from confocal z-stack of fluorescent cores in colloidal crystal close to the cover glass, recorded with a Leica HC PL APO 100× /1.40 OIL STED WHITE objective (2 pixel 3D median filter), showing the response function of the microscope in the lateral plane (left), *XZ* plane (middle) and *YZ* plane. The two axial planes are marked by the dashed lines in the lateral plane (left). Alignment of 2D (b) and 3D (c,d) STED excitation (cyan) and depletion (magenta) lasers in the lateral plane, as imaged using a Leica HC PL APO 93× /1.30 GLYC motCORR STED WHITE objective in reflection mode (left) using a gold core particle close to (b,c) and 50 µm (d) from the cover glass, and the resulting STED images (2 pixel median filter) of the fluorescent cores in the crystal (right), showing increased resolvability of the particles in the crystal as compared to a). Pinhole size in fluorescence images: (a) 0.7 and (b-d) 1 Airy units, in all reflection mode images is 4.7 Airy units. All scale bars are 500 nm.

The gold cores in the crystal were used to check the shape of the depletion laser spot and the overlap with the excitation laser spot, which are both critical for an optimal STED resolving power. Figure 3b shows the excitation spot (cyan) and the 2D STED ^31^ ‘doughnut’-shaped depletion spot (magenta) imaged in reflection mode using a Leica HC PL APO 93×/1.30 GLYC motCORR STED WHITE objective with the pinhole set to 4.7 Airy units. In 2D STED the PSF is only improved in the *XY* plane, contrary to 3D STED where the PSF is also modified along the optical axis in the *Z* direction ^32^. Subsequent imaging of the fluorescent cores with the pinhole set to 1 Airy unit shows an increase in lateral resolution of the STED microscope (Figure 3b) as compared to confocal (Figure 3a), even when using a lower NA objective.

Since 2D STED microscopy only improves the resolution in the lateral plane compared to confocal microscopy, it is primarily employed on thin samples, close to the cover glass. 3D STED, however, enables imaging at sub-diffraction resolution in both lateral and axial directions, which allows imaging of samples at sub-diffraction resolutions also far away from the cover glass ^32^. Figure 3c-d shows the excitation and depletion laser spots of a 3D STED confocal microscope equipped with a Leica HC PL APO 93×/1.30 GLYC motCORR STED WHITE objective as imaged in reflection mode close to the cover glass (Fig. 3c) and ∼50 µm away from the cover glass (Fig. 3d). The correction collar (CC) of the objective was used to keep the intensity of the top and bottom depletion spots balanced when imaging more than ∼30 µm from the cover glass. To check the results of the alignment of the lasers, the fluorescent cores were imaged in 3D STED imaging mode at a pinhole size of 1 Airy unit (Figure 3c,d). The improved axial resolution of 3D STED imaging is visible from the decreased size of the PSF in the *Z* direction (FWHM ≈ 300 nm) as compared to the confocal PSF (FWHM ≈ 600 nm, Figure 3a), even at ∼50 µm away from the cover glass.

The presented sample enables quality control of the PSF, but the measurement of its extended structure requires a greater separation between the fluorescent beads. An increased separation allows imaging of a single bead, without its axial lobes overlapping with those of neighboring particles. Therefore, a slightly modified sample was constructed. This consisted of the same probe particles as described above, arranged not in a dense crystal but sparsely distributed in 3D across a scaffold of unlabeled silica spheres (310 nm, 10% PDI, Supplementary Fig. 2), embedded in a refractive index matching solvent mixture. While shown here in a separate sample, this can also be realized by incorporating sparse spheres with a fluorescently labeled core in the aforementioned crystal (shown in Fig. 1), where the dye of the sparse spheres differs from the dye of the fluorescent particles in the crystal.

Even the slight refractive index mismatch between the immersion and sample media (in this case: 1.45 vs. 1.42) causes increased aberration at increasing imaging depth. This aberration was minimized by attempting to optimize the position of the objective CC using images of the dilute gold-core particles. Two strategies to find this ‘optimal’ position were tested: maximize the intensity of the excitation reflection (cyan in Figure 4) or equalize the intensity of the axial 3D STED lobes (magenta in Figure 4) which become lopsided deep in the sample without CC adjustment (Supplementary Figure 3). The two strategies were found not to agree on an optimal position, so the latter was chosen to ensure optimal STED imaging. The equalized lobes at each height are shown in the top row of Figure 4.

**Figure 4:**
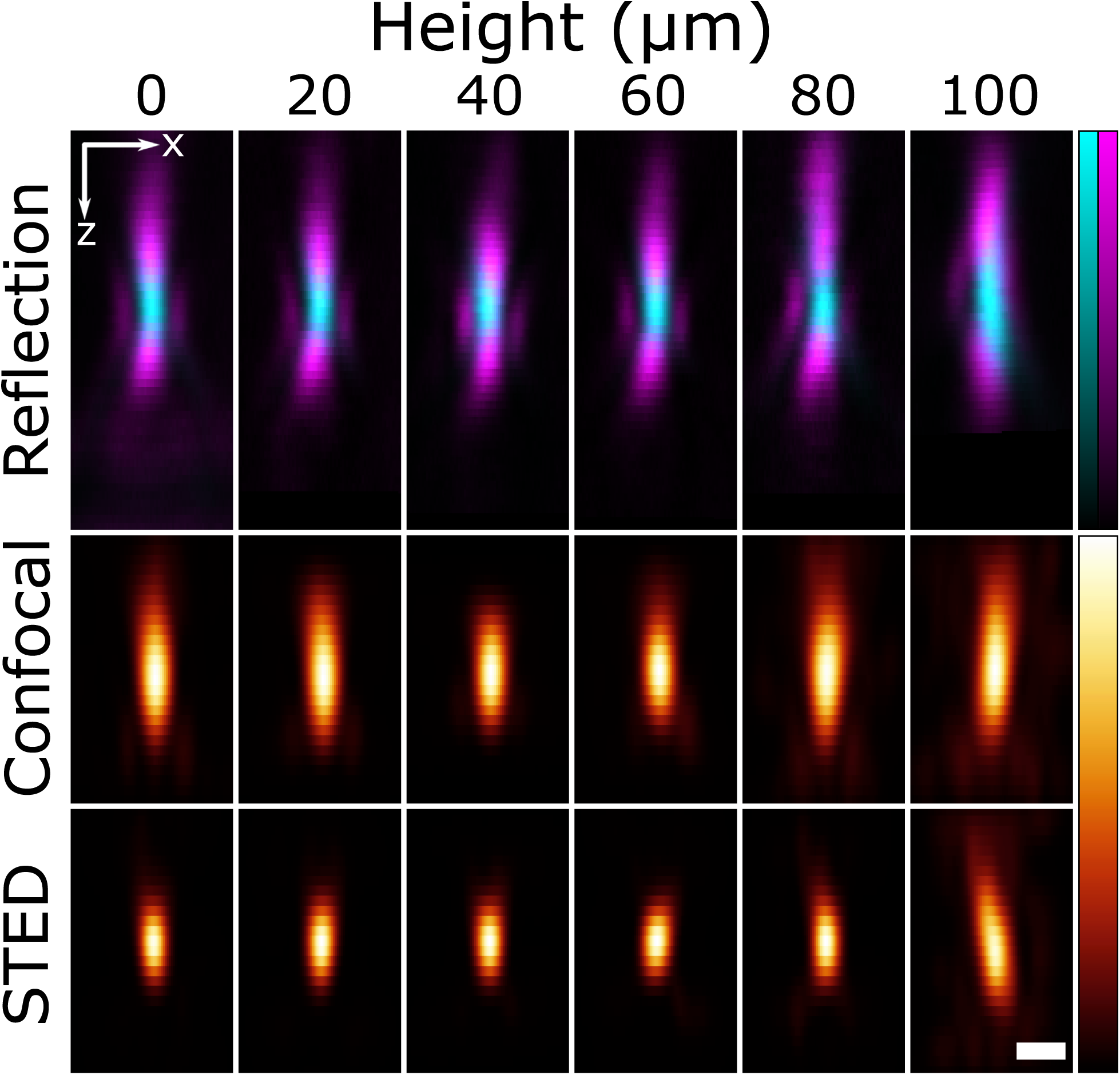
Confocal and 3D STED point spread functions at different distances from the cover slip. The excitation (cyan) and depletion (magenta) lasers imaged in reflection using the sparse gold-core particles (top row). To correct for the mismatch between the immersion liquid of the objective and the sample, at every height the correction collar (CC) of the objective was adjusted to equalize the peak intensities of the axial STED lobes (see also Supplementary Fig. 3). Point spread functions in the *XZ* plane measured in confocal (middle row) and 3D STED (bottom row) mode using silica coated fluorescent beads dispersed in a disordered scaffold of unlabeled silica spheres (310 nm, 10% PDI, Supplementary Fig. 2). At least 10 well separated beads were recorded and averaged at each height, and the PSFs were extracted by deconvolution with a sphere the size of the fluorescent cores. The scale bar is 500 nm.

By imaging 3D volumes at various heights above the coverslip (using the previously determined CC positions) in both confocal and 3D STED imaging modes, the PSFs shown in Figure 4 (middle and bottom row) were obtained. Several (*>* 10) beads at each height were averaged and the PSF at that height was computed by deconvolution with a sphere the size of the fluorescent core^14^. The ability of measuring PSFs far from the cover glass was demonstrated using beads at depths up to 100 µm. This enables quality control of the microscope, with this particular setup being found to maintain a compact 3D STED PSF at depths of up to 80 µm. Furthermore, measurements of the PSFs at different heights enables accurate deconvolution and high precision analysis of images captured deep inside thick specimens^8–10^.

Finally, to show the importance of a proper alignment of the excitation and depletion spots in STED microscopy, fluorescently labeled *α*-tubulin in HeLa-cells was imaged using 2D STED microscopy using a Leica HC PL APO 100×/1.40 OIL STED WHITE objective (Figure 5). When the two laser spots were properly aligned using our sample, an increase of the lateral resolution is visible (Figure 5a-d). When the minimum of the depletion spot is slightly off-center with respect to the maximum of the excitation spot, the depletion of the fluorophores by the depletion beam results in a higher level of noise in the resulting image and no improvement in the resolution compared to the confocal image (Figure 5f-h).

**Figure 5:**
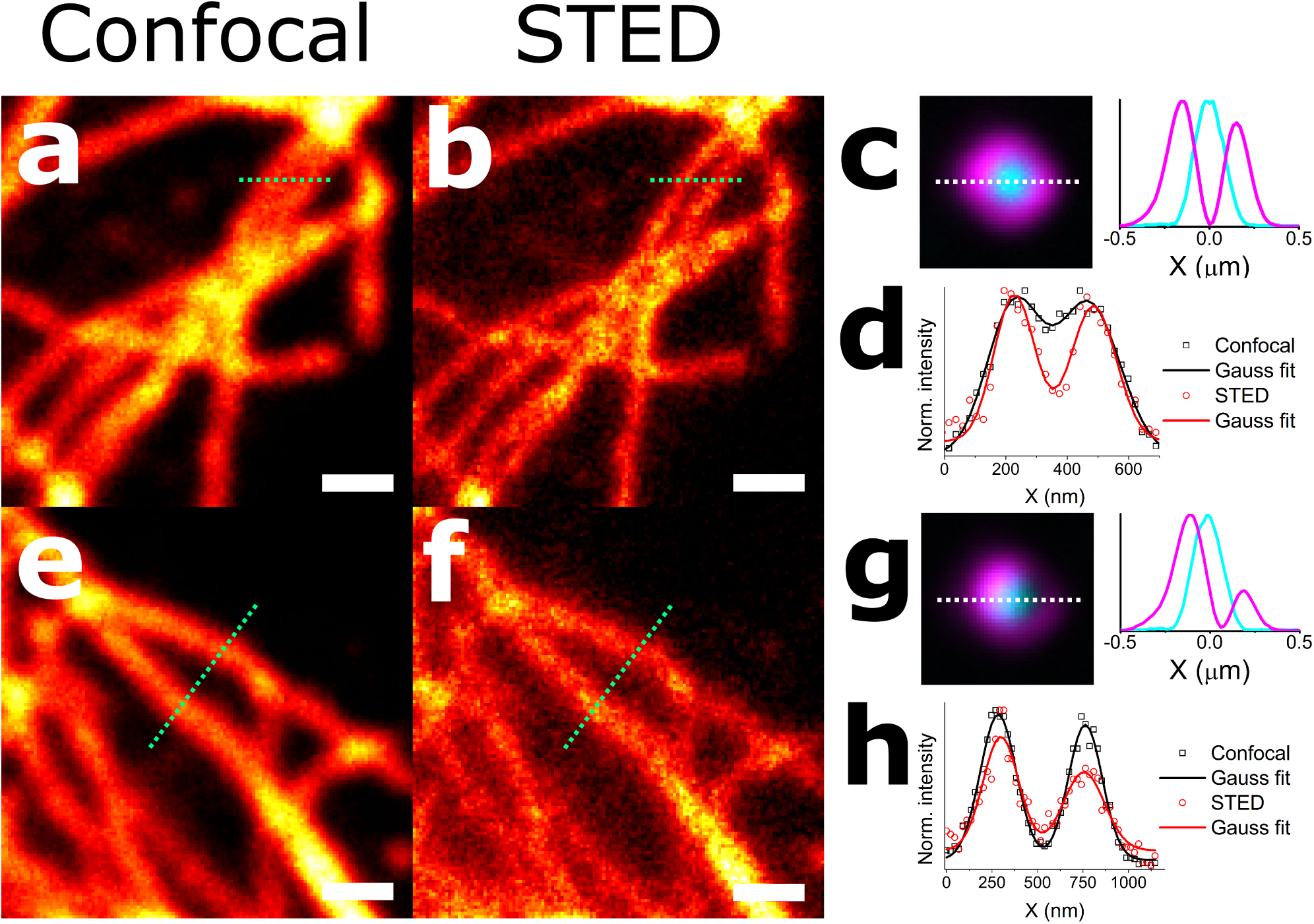
The importance of a proper alignment of the STED microscope. Results of alignment of the laser beams in STED microscopy on imaging of *α*-tubulin (HeLa-cell) with a HC PL APO 100× /1.40 OIL STED WHITE objective. a) Confocal micrograph. b) 2D STED image at same region as a) after aligning the minimum of the ‘donut’-shaped depletion laser spot to the maximum of the excitation laser spot. c) Alignment of excitation (cyan) and STED (magenta) lasers in b). d) Intensity profiles along the lines in a) and b), showing the increased resolving power of the STED compared to confocal microscopy. e) Confocal micrograph. f) 2D STED image at same region as e) with slightly misaligned excitation and depletion laser spots. g) Alignment of excitation (cyan) and STED (magenta) lasers in f). h) normalized intensity profiles along the lines in e) and f), showing no increase in the resolving power of the (2D) STED compared to confocal microscopy. Scale bars are 500 nm.

## Discussion

The sample as presented here is a robust standard for the quality control of a laboratory’s confocal and STED microscopes. By the time of writing the shelf life of the sample is on the order of months. Although the small fluorescent cores tend to bleach upon prolonged excitation the sample can be used repeatedly because the crystal contains over 10^10^ particles due to its larger size (close to 2 cm^2^, see Fig. 1c).

In the measurement of the characteristic lateral length scales Δ_*X*_ and Δ_*Y*_ an error below 1% is found (obtained by measuring at three depths). In addition, for the axial interlayer spacing an error of less than 0.5% is found (obtained from a linear fit of the intensity peaks of the crystal layers). This low error is supported by the perfect linearity of the axial interlayer spacing as found by FIB-SEM tomography and particle tracking. These low errors make the sample suitable for the calibration of microscopes at a high precision.

Due to the bottom-up assembly, the sample is highly tunable. This makes it possible to adjust the sample to, for example, meet the requirements for different high NA lenses (e.g. oil immersion) and/or different techniques. This can be achieved by changing the core, the ratio between different types of cores, the refractive index of the particles, the shell and the total size of the particles, which will be discussed next.

While it was demonstrated that rhodamine B labeled silica cores can be used, the synthesis method of these cores allows for the incorporation of a wide variety of dyes^33–39^. This can be useful because by mixing cores with different dyes in the non-fluorescent silica growth step, one can accommodate microscope setups with multiple laser lines. Instead of keeping the gold and dye separated within the sample, the two can also be combined within a single particle, by growing a fluorescent layer around the gold core. Another option is to use a silver core instead of a gold core. This flexibility opens up the possibility to design nanocomposite particles where the fluorescence is enhanced by the presence of the metal core^40^, which has recently been demonstrated to be compatible with STED microscopy^41^.

The possibility to exchange the fluorescent cores with quantum dots is also worthwhile exploring, since quantum dots have excellent photostability, are only a few nanometer in size and have shown to be compatible with STED microscopy^20, 21^. As the absorption and emission wavelengths of quantum dots are size dependent^42^, they can be tailored to the wavelength of the excitation and STED lasers. When measuring a PSF using quantum dots, the challenge is to ensure there is enough separation between them which can be achieved by coating the quantum dots with silica^43–45^. This seeded growth method is directly compatible with the work presented here, because silica coating is performed in the same reverse micro-emulsion system used to synthesize the fluorescent cores. The most promising quantum dots to use are so-called CdS/CdSe/CdS quantum well dots, as these can be coated with silica while maintaining a high photoluminescence quantum yield^46, 47^.

Next to quantum dots, a variety of nanometer sized probes have been put to use in STED imaging^48^, such as fluorescent nanodiamonds^49^, upconversion nanocrystals^50^ and carbon dots^51^. Although these probes can be coated with silica^52–54^, and could be integrated in our demonstrated sample, no STED microscopy has been demonstrated on these probes after silica coating.

Another method to tune the sample is by tuning the silica shell that is used to ensure separation between the cores. While in this work the focus was on a refractive index of 1.43 compatible with glycerol objectives, it can also be tuned to a higher refractive index. This can be achieved by synthesizing a shell of titania/silica composite with a higher refractive index^55–57^. Doing so, the refractive index can easily be increased to match the refractive index of oil objectives (1.52), or any other intermediate refractive index that is comparable to the refractive index of life science specimens of interest. Another option is to increase the size of the shell to increase the spacing between the probes. This makes the sample compatible with lower NA objectives, where the axial resolving power is smaller. For this, the assembly method needs to be adjusted due to a faster settling rate of the particles, as has been demonstrated for close to 1 µm silica particles^58^.

Finally, by introducing a refractive index mismatch between the particles and the embedding solvent, the effect of the mismatch on the PSF can be measured as a function of the distance to the cover glass. Moreover, the periodicity of the crystal in the axial direction and the known refractive indices of the beads and the embedding mixture can be used to test effective medium theories developed to determine an effective refractive index in optically heterogeneous environments ^59^.

## Methods

### Particle synthesis: silica-coated gold cores

Gold nanoparticles (80 nm, in citrate buffer (OD 1), Sigma-Aldrich) were functionalized with polyvinylpyrrolidone (PVP, Mw = 10,000 g mol^−1^) by transferring 20 mL of the gold nanoparticle solution and 400 µL of a 10% (w/v) PVP solution (1 g in 10 mL water) to a vial ^60^. The obtained solution was stirred for 16 hours and was transferred to 5 mL eppendorf tubes and centrifuged 5 minutes at 5.000 rcf. The supernatant was removed as much as possible with a glass pipette and the gold nanoparticles were redispersed in 2.5 mL ethanol and collected in a 4 mL glass vial.

Silica coating was performed in a closed 4 mL vial under constant stirring at 600 rpm and was initiated by the addition of 250 µL ammonia (28-30 *w*% NH_3_ basis, ACS reagent, Sigma-Aldrich) and 25 µL of a 10 vol% solution of tetraethyl orthosilica (TEOS, reagent grade, 98%) in ethanol (absolute, Merck) ^60^. These additions were followed by the addition of 50, 100 and 200 µL of 10 vol% after 90, 270 and 360 minutes. 90 minutes after the final addition, the solution was transferred to a 20 mL vial and diluted with ethanol to obtain a total volume of 20 mL. This diluted solution was centrifuged 15 minutes at 1.000 rcf and the white turbid supernatant was removed. Next, the particles were redispersed in 20 mL ethanol by sonication. Centrifugation and redispersion of the particles was repeated to remove empty silica spheres formed by secondary nucleation. After the final centrifugation step, the particles were redispersed in 10 mL ethanol. The resulting particles and their gold cores were imaged with transmission electron microscopy (TEM) (Supplementary Fig. 4), from which an average particle diameter of 321.5 nm (4.3% polydispersity index (PDI), where PDI is the standard deviation over the average diameter) was determined by measuring 100 particles.

### Particle synthesis: silica-coated fluorescent cores

Dye-APTES coupling was performed by transferring 6.00 mg Rhodamine B isothiocyanate (mixed isomers, RITC, Sigma-Aldrich), 500 µL ethanol and 12 µL (3-Aminopropyl)triethoxysilane (APTES, 99%, Sigma-Aldrich) to a vial ^38,39^. The vial was wrapped in aluminium foil and stirred for 5 hours. A reverse micro-emulsion was prepared by transferring 50 mL cyclohexane (Sigma-Aldrich) and 6.5 mL Igepal CO-520 (Sigma-Aldrich) to a vial under vigorous stirring (700 rpm) ^61–63^. As soon as a clear solution was obtained, 400 µL TEOS was added to the solution. After 2 minutes stirring, 50 µL of dye-APTES solution was added resulting in the formation of a colorless solution. After an additional 5 minutes of stirring to ensure complete homogenisation, 750 µL ammonia was added to initiate the reaction. Immediately after this addition, the solution turned pink. After an additional minute of stirring the solution was stored in a dark place for the reaction to proceed. After 24 hours, the reaction mixture was transferred to a round bottomed flask. The cyclohexane was evaporated under reduced pressure (∼10 mbar) using a rotary evaporator. The flask was placed in a room temperature water bath and evaporation was performed under vacuum. After 20 minutes, a very viscous, pink solution was obtained of fluorescent silica particles dispersed in the non-ionic surfactant. 10 mL *N,N*-Dimethylformamide (DMF, Sigma-Aldrich) and 10 mL ethanol were added to this liquid resulting in the formation of a clear, non-scattering, pink solution.

Further silica growth of the fluorescent cores was performed by transferring 5.00 mL of the fluorescent core solution, 3.45 mL ethanol, 1.11 mL water and 0.45 mL ammonia to a three necked round bottomed flask ^64–66^. Under gentle stirring (200 rpm) and nitrogen flow a 3 times diluted solution of TEOS in ethanol was added to the solution using a syringe pump. A water/ammonia/ethanol solution was added simultaneously to keep the water and ammonia concentrations constant. A total of 6.54 mL was added at a flow rate of 0.32 mL/h, after which the flow rate was doubled. After 10.98 mL of TEOS solution was added, the syringes were refilled and TEOS addition was continued until a total volume of 30.13 mL was added. After silica growth, the solution was transferred to two 50 mL eppendorf tubes and centrifuged 30 minutes at 2000 rcf. Particles were redispersed in 20 mL absolute ethanol by sonication, collected in one vial and stored in a fridge at ∼4 ^*°C*^. The resulting particles and their fluorescent cores were imaged using TEM (Supplementary Fig. 4) and an average diameter of the cores and silica-coated particles were 44.7±2.3 and 240.1± 4.1 nm, both obtained by measuring the diameter of 100 particles. To match the size of the gold-core silica particles, the fluorescent-core silica particles were grown further. 93 mL ethanol, 1.80 mL of silica coated fluorescent cores and 12.4 mL water were transferred to a 250 mL round bottomed flask. The obtained solution was sonicated for an hour before growth was started. Under gentle stirring (200 rpm) and nitrogen flow, 3 mL of a 3 times diluted solution of TEOS in ethanol was added to the solution using a syringe pump with a flow rate of 0.65 mL/h. A water/ammonia/ethanol solution was added simultaneously to keep the water and ammonia concentrations constant. After this first growth step, a TEM sample was prepared to determine whether the size of the particles matched the size of the silica coated gold particles.

### Particle synthesis: further growth of a mixture of gold- and fluorescent-core particles to 500 nm diameter particles

Further growth of a mixture of gold- and fluorescent-core silica particles was performed to obtain particles with a total diameter of approximately 500 nm ^64–66^. The gold- and fluorescent-core silica particles were mixed in a 1 to 100 number ratio in ethanol, by adding 8.8 mL of the suspension of silica coated gold particles to the reaction mixture containing the fluorescent-core silica particles (see previous paragraph). The concentration of gold particles and the weight fraction of the fluorescent cores were used here to determine how much of the gold-core suspension should be added to obtain the desired 1:100 ratio between the two types of particles. Next, growth was continued by the simultaneous addition of 17.16 mL of the 3 times diluted solution of TEOS in ethanol and the water/ammonia/ethanol solution with a flow rate of 1.00 mL/h. After silica growth, the solution was transferred to three 50 mL eppendorf tubes and centrifuged 30 minutes at 500 rcf. After repeated centrifugation and redispersion of the particles, all particles were redispersed in 40 mL absolute ethanol by sonication, collected in one vial and stored in the fridge. The resulting particle mixture was imaged using TEM (Supplementary Fig. 6), from which the average diameter was determined: 505.1 nm (1.8 % PDI), by measuring 368 particles.

### Particle synthesis: non-fluorescent silica particles for scaffolding

15 mL ammonia and 120 g ethanol were mixed in a 250 mL round bottomed flask and the temperature was raised to 30 ^*°C*^ and stirred using a stir bar ^64,67^. To this mixture, 6.60 mL TEOS was pipetted under the liquid surface with vigorous stirring at 600 rpm. After 9 minutes, blue scattering of the particles was observed and the stirring speed was reduced to 300 rpm. 22 h after the TEOS addition, the mixture was poured into a 500 mL centrifuge bottle, ethanol was added and the mixture was centrifuged at 700g for 45 minutes. The supernatant was replaced with ethanol and this washing procedure was repeated twice. This resulted in polydisperse particles with a diameter of 310 nm (10% PDI, Supplementary Fig. 2c).

### Colloidal crystal growth

Colloidal crystal growth was performed via the vertical deposition method^24^ at elevated temperature to speed up the evaporation process^68–70^. Briefly, 8.0 mL of a 1 vol% mixture of gold- and fluorescent-core particles in ethanol with a diameter of ∼500 nm was transferred to a 20 mL glass vial. A cover glass (Marienfeld Superior #1.5H, 24×50 mm) was placed upright inside the solution under a small angle (∼5^*°*^) with respect to gravity. This vial, and a 100 mL beaker filled with ethanol were placed inside a 50 ^*°C*^ preheated oven (RS-IF-203 Incufridge, Revolutionary Science) and covered with a large beaker (upside down). After approximately 16 hours, the cover glass was removed from the solution. An opaque deposition of particles was observed on the cover glass. Any particles sticking to the back of the cover glass were removed with an ethanol soaked paper tissue. Bragg reflections were observed which indicates the formation of a crystalline structure. For the growth of thicker crystals, growth steps were performed by repeating the vertical deposition method up to three times.

### Calibration and alignment sample preparation

Two spacers (Menzel #00 cover glass, thickness = 55–80 µm) were glued onto a microscope slide using UV-glue (Norland 68 Optical Adhesive), about 5 mm apart. Next, the cover glass containing the crystal was glued onto the spacers, such that the crystal was inside the created channel. This channel was then infiltrated with a mixture of *n*-butanol and glycerol (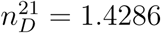, measured using an Atago 3T refractometer) in order to refractive index match the particles to the surrounding liquid, to reduce scattering and optimize the imaging conditions. The use of water in the matching solvent mixture was omitted, as it has been reported to change the refractive index of silica particles ^71^, because it is small enough to enter the interior of the silica network, while molecules larger than 0.3 nm cannot. Finally, the channel was closed using UV-glue. While curing the glue, the crystal was covered with aluminium foil as protection from the UV radiation to prevent bleaching of the dye in the particles.

### HeLa cell sample

A HeLa cell sample (Active Motif/Leica Microsystems) was used for evaluation of the alignment of 2D STED imaging of a realistic biological sample, where the *α*-tubulin was stained with BD Horizon V500-streptavidin (BD Biosciences) for imaging. The cells in the sample were mounted using Mowiol 4-88 on a 170 ± 10 µm thick cover glass (Hecht Assistent #1014), resulting in a sample refractive index of *n* ≈ 1.52.

### Confocal and STED measurements

Calibration and alignment measurements were performed on an inverted Leica TCS SP8 STED 3X confocal microscope, equipped with a Leica HC PL APO 100×/1.40 OIL STED WHITE or a Leica HC PL APO 93×/1.30 GLYC motCORR STED WHITE objective with a correction collar (CC). The confocal microscope, as well as the CC were controlled with LAS X software (version 3.5, Leica Microsystems). In case of the oil objective Type F immersion oil (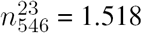, Leica Microsystems) was used as immersion liquid, while for the glycerol objective a 85 w% glycerol/water mixture 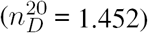 was used. To image the fluorescent cores of the particles, a pulsed (80 MHz) super-continuum white light laser (SuperK, NKT Photonics) was tuned at 543 nm, while the emission was detected using a gated (0.3-6.0 ns) HyD detector (553-650 nm). For STED imaging of the fluorescent beads a continuous wave (CW) depletion laser with a wavelength of 660 nm was used. The depletion laser was either fully focused into a ‘doughnut’-shaped spot (2D STED ^31^) or into an axial depletion spot (3D STED ^32^). The gold cores were imaged by detecting the reflection of the excitation or depletion laser using a photomultiplier tube (PMT). The CC of the glycerol objective was used to compensate for the refractive index mismatch between the sample (1.43) and the objective (1.45), and allowed to fine tune the depletion laser pattern, as discussed in more detail in the supplementary information of [22].

For the *α*-tubulin images, the white light laser was tuned to 470 nm, with the wavelength range of the HyD detector set to 485-580 nm while a 592 nm CW depletion laser was used. Acquisition parameters of the confocal and STED images are listed in Supplementary Table 1. Any image filtering was done using Fiji ^72^ (ImageJ 1.52d).

### Image deconvolution

The confocal 3D stack of the fluorescent beads (Figure 2) was deconvolved with a theoretical point spread function using Huygens Software (Scientific Volume Imaging, 17.04).

### Calibration measurements

For the lateral calibration Fast Fourier transforms of the XY slices were calculated using iTEM (Soft Imaging System GmbH, 5.0) and a 1 pixel Gaussian blur was applied using Fiji^72^ to reduce noise. To measure the Δ_*X*_ and Δ_*Y*_ distances (see Fig. 2e) intensity profiles parallel to the direction of interest were drawn through the origin. Using OriginPro (v8.0891, OriginLab Corporation) Gaussian functions were fitted to the peaks nearest to the origin in the intensity profiles, to obtain the distance of these peaks to the origin Δ_*X*_ and Δ_*Y*_.

For the axial calibration Fiji was used to plot the intensity of the deconvolved confocal z-stack as a function of *Z*-height. Using OriginPro the peaks in the profile were fitted with Gaussian functions. The peak positions were plotted as a function of peak number and fitted using a linear function using OriginLab. The slope of this function was used as interlayer distance.

### Measuring the scaling factor of a 93×/1.3 NA glycerol objective for n=1.43 samples

To determine the scaling factor of the axial distances in our sample as measured by confocal microscopy, we used the method described in [14]. A sample cell was constructed by gluing two Menzel #00 cover glasses as spacers on a microscope slide, after which a channel was created by bridging the spacers with a Menzel #1.5 cover glass. The Fabry-Pérot fringes were measured using a Fourier-transform infra-red (FTIR) spectrometer (Vertex 70, Bruker) and fitted to determine the cell height: 94.257±0.008 µm. Next, rhodamine B dyed poly(methyl methacrylate) spheres (70 nm diameter) were deposited from hexane on the inside of the cell and the height of the cell was measured in fluorescent mode on a confocal microscope equipped with a 20×/0.7 NA air objective: 98.0±0.2 µm, resulting in a miscalibration of the microscope stage of 4.0±0.2%. After filling the cell with the glycerol/*n*-butanol mixture (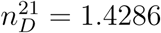) and closing it with UV-glue a height of 99.9±0.2 µm was measured, which was corrected for the miscalibration of the microscope stage: 96.0±0.3 µm. This gives a scaling factor of 0.982±0.003 for axial distances in a sample with a refractive index of 1.43 recorded with a 1.3 NA glycerol objective.

### PSF measurement from sparse probe particles

The probe particles in Figure 1a were distributed in 3D across a scaffold of undyed silica particles. This was achieved by mixing dispersions of the particles in ethanol such that the number ratio of scaffold : fluorescent : gold-core particles was 85000 : 99 : 1. Next, 5 µL of this 18 mg/mL dispersion was repeatedly dropcast on the same spot on a #1.5H cover glass, rapidly evaporating the ethanol in an oven at 50 ^*°C*^ between each drop addition. The resulting residue was uneven with regions more than 100 µm high after several steps. This residue was incorporated into a microscopy cell of the type shown in Figure 1a (see also Supplementary Fig. 2). The refractive index of the scaffold particles was measured using the method described in Supplementary Note 2 and the cell infiltrated with a liquid mixture of the same refractive index, specifically 63.0 w% glycerol/water 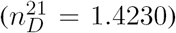, before sealing with UV-glue.

Before imaging the fluorescent particles, the reflection signals from the gold core particles were used to find optimal parameters for 3D STED imaging. First, the STED alignment was adjusted by centring the excitation and depletion reflection patterns (cyan and magenta respectively in Figure 4) of a gold-core particle located on the cover glass. Then, a CC position was chosen to equalize the 3D STED axial lobes as viewed in the *XZ* plane through the centre of the particle. This was achieved by iteratively adjusting the CC position, re-centring the particle in the field of view by moving the sample stage and checking the intensities of the side lobes with a line profile along *Z* through the *XY* centre of the particle. With these equalized as much as possible, the reflection images shown in the top row of Figure 4 were captured, using the parameters and CC positions in Supplementary Tables 1 and 2, respectively.

Next, image volumes of the fluorescent beads were captured from which to extract the PSFs. At each height, test images using low laser power were captured at different *XY* positions to discern whether or not any fluorescent beads were located in the region of interest, away from the edges. If so, the position was marked and eventually 7–15 *XYZ* stacks of dimension 10.4 × 10.4 × 4.1 µm were captured at the various chosen *XY* positions in a ‘Mark and Find’ acquisition. The checking was required to avoid wasted image volumes containing no usable particles and it was done in this way to avoid bleaching the fluorescent cores before the main measurement. The low pixel dwell time (0.3 µs) used was to ensure that the beads did not bleach significantly during acquisition. Approximately thirteen image volumes were recorded at each height, each with imaging times of around 1 minute. The confocal PSFs were recorded first, followed by the STED ones, with 5 hours between the first and last measurements. The room temperature was maintained at 23.5 ^*°C*^ throughout this period.

The beads in the image volumes at each height were located and summed together to give an averaged image of the particles at that height. Only signals well separated from each other and away from the image volume boundaries were included. This average at each height was deconvolved with a sphere of diameter 45 nm (the known fluorescent core size), yielding the measured PSF^73^. Both of these steps were carried out using Huygens Professional deconvolution software (version 17.04, SVI).

## Acknowledgements

The authors are grateful to Dave van den Heuvel for the help with confocal measurements and particle synthesis, Peter Helfferich for the help with confocal (STED) measurements, Michiel Hermes for the particle identification in the FIB-SEM data and Job Fermie for the help with FIB-SEM sample preparation. This project was funded by the European Research Council under the European Union’s Seventh Framework Programme (FP-2007-2013)/ERC Advanced Grant Agreement 291667 HierarSACol, NWO – Microscopy Valley under project number 12715 and the Netherlands Center for Multiscale Catalytic Energy Conversion (MCEC), a NWO Gravitation programme funded by the Ministry of Education, Culture and Science of the government of the Netherlands.

## Author Contributions

AvB initiated the project. HG and AvB supervized the research. JF, CLK and MdPP synthesized the particles. JF, CLK, MdPP and EBvdW performed the refractive index measurements. JF determined fluorophore labeling density of the particles. JF, MdPP, EBvdW and PNAS prepared the colloidal crystals. EBvdW, CLK, MdPP and PNAS performed the confocal (STED) microscopy measurements. DAMdW performed the FIB-SEM measurements. EBvdW, JF, HG and AvB co-wrote the manuscript. All authors discussed the results and commented on the manuscript.

## Competing Interests

The authors declare that they have no competing financial interests.

## Notes

### Competing Interest Statement

The authors have declared no competing interest.

