## Supplementary Information for "3D test sample for the calibration and quality control of super-resolution and confocal microscopes"

### 1 Supplementary Figures

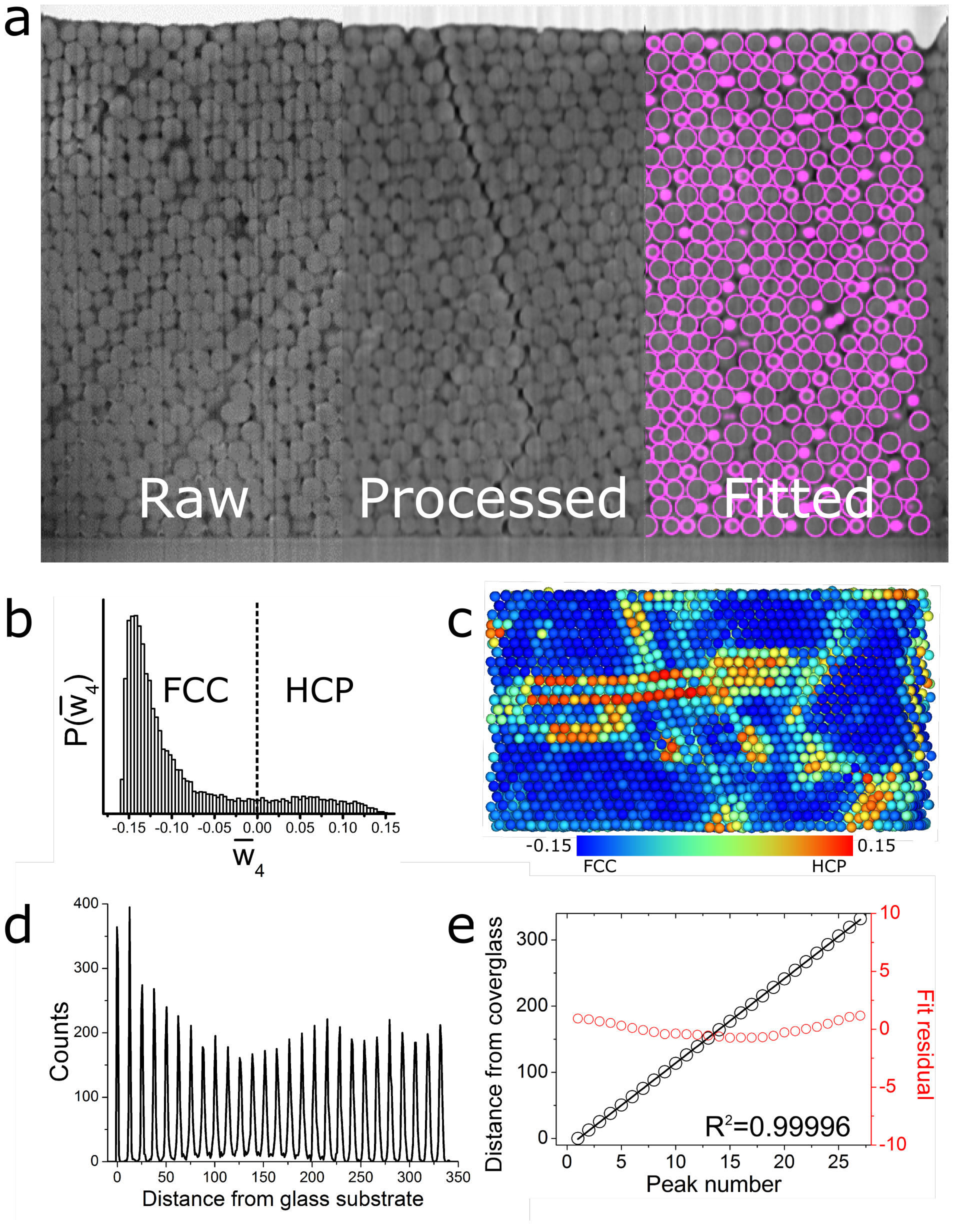


Supplementary Figure 1: **FIB-SEM tomogram of colloid crystal.** a) Slice from FIB-SEM tomogram. As recorded (left), after filtering (middle) and with overlay of fitted particle coordinates (right). b) Histogram of the $\overline{w}_4$ bond orientational order parameter values [1] calculated from the coordinates in a), confirming the face-centered cubic structure of the crystal. c) Computer rendering of the particles from coordinates obtained by particle identification, coloured to the $\overline{w}_4$ values of the particles. d) Histogram of distance from the cover glass for every coordinate demonstrating the layering of the particles in the crystal. e) Linear fit of the peak positions in d) confirming close to perfect linearity.

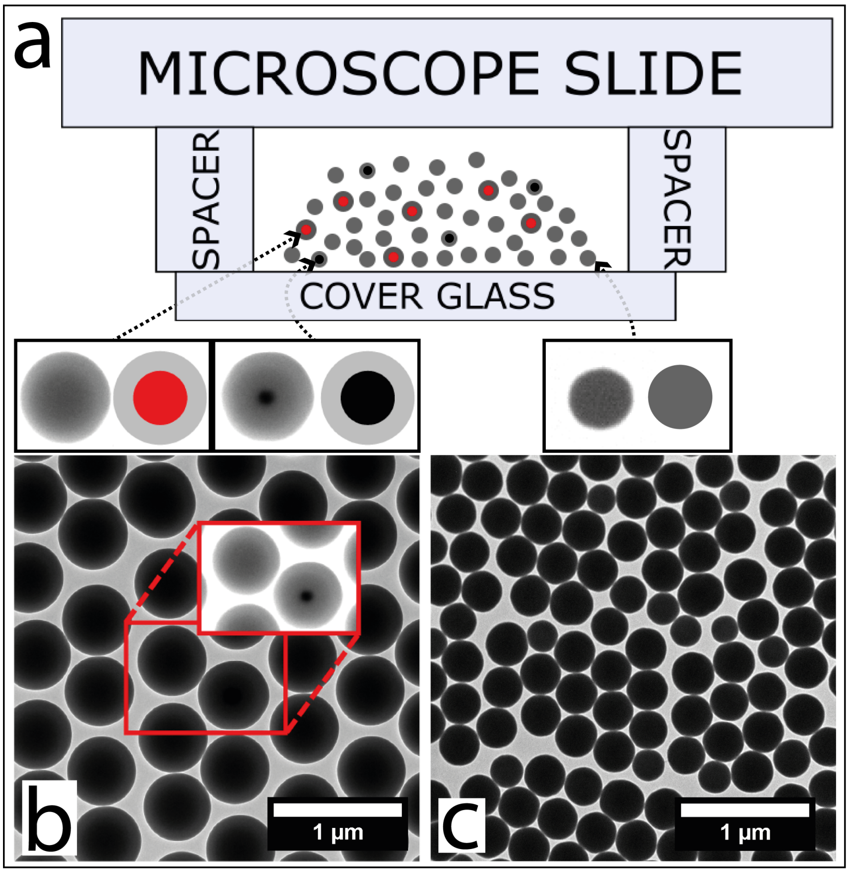


Supplementary Figure 2: Ternary particle system used for PSF measurements. a) Schematic of the cell showing the fluorescent and gold-core tracer particles (red and black centers respectively) suspended in a majority of 'undyed' (host) particles (no heterogeneous core). The number ratio of host:fluorescent:gold-core tracer particles was $85000 : 99 : 1$ resulting in a mean separation of $\sqrt[3]{850/0.99} \approx 10$ scaffold particles between fluorescent beads, or around 3 μm. b) TEM image of the mixture of fluorescent and gold core particles. The increased contrast in the inset allows the gold core to be seen. c) TEM image of the polydisperse scaffold particles at the same scale.

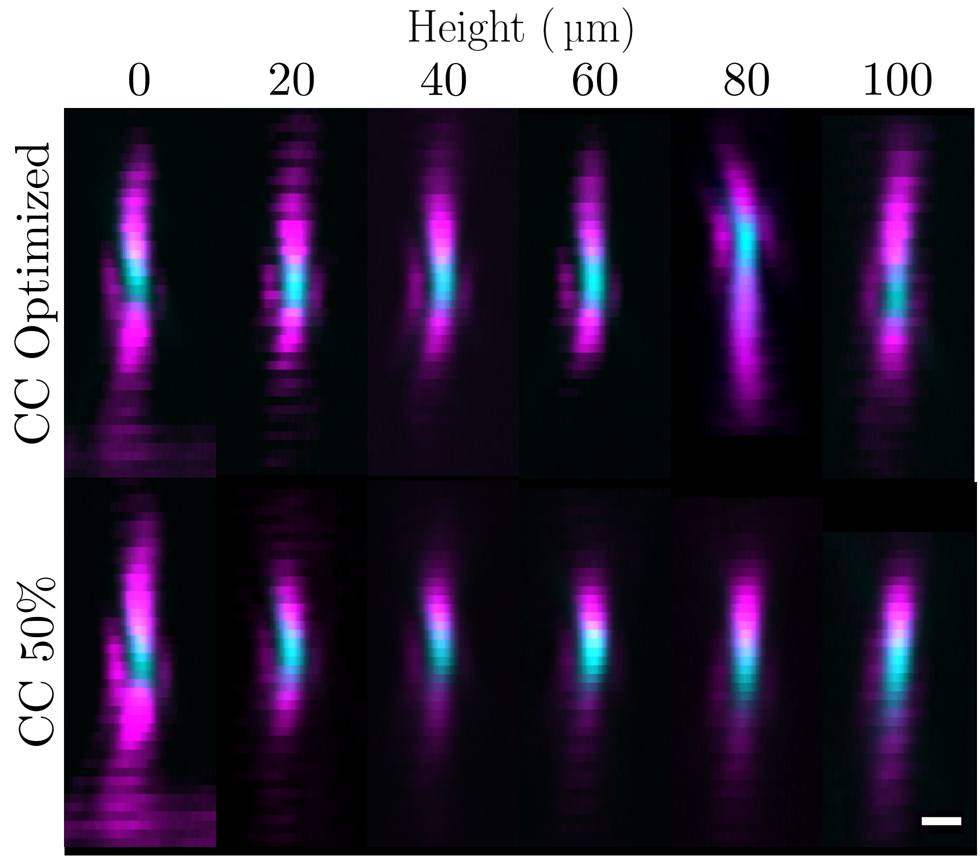


Supplementary Figure 3: Effect of the objective correction collar on the depletion beam. The top row shows reflection of the excitation (cyan) and depletion (magenta) beams at different heights with the correction collar optimized (see Table 1). The bottom row shows the same with the correction collar position fixed at 50%. The scale bar indicates 500 nm.

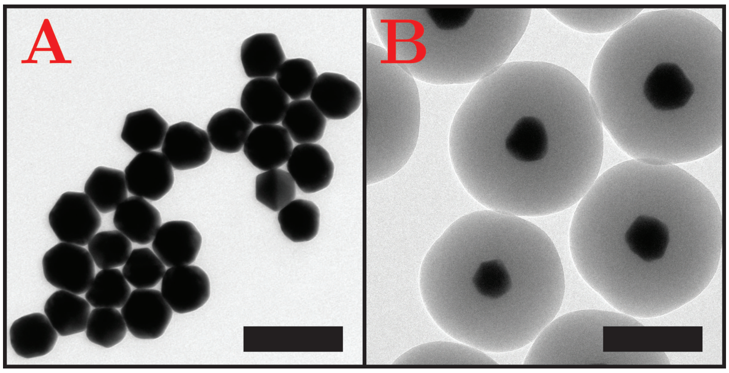


Supplementary Figure 4: Representative TEM images of the 80 nm gold nanoparticle before (A) and after (B) silica coating. The scale bars represent 200 nm.

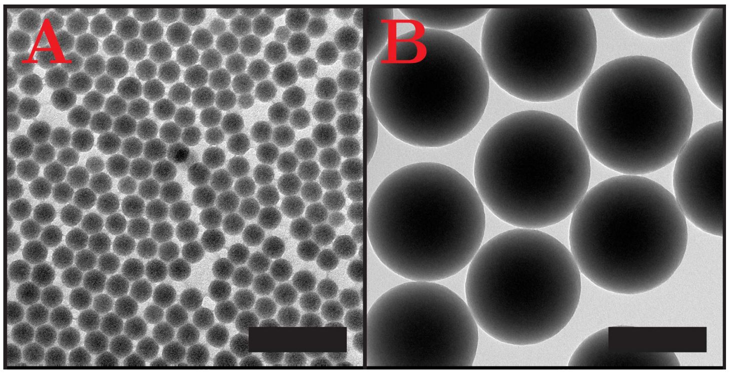


Supplementary Figure 5: Representative TEM images of samples of the fluorescent cores before (A) and after (B) further silica coating. The scale bars represent 200 nm.

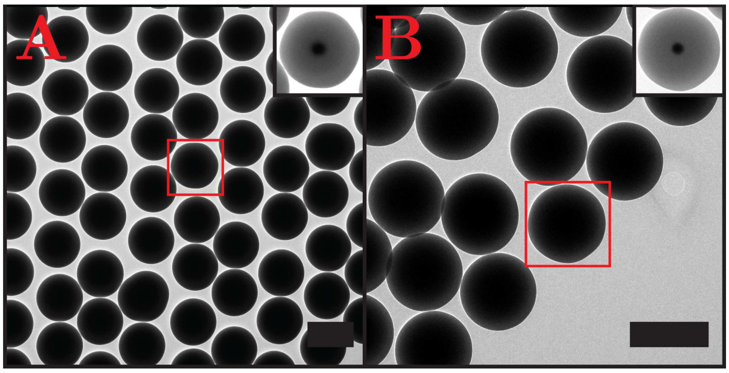


Supplementary Figure 6: Representative TEM images after the final silica growth step to ∼500 nm of the mixture of the two types of core-shell particles (gold and fluorescently labeled silica cores). In the inset of (A) and (B) the contrast of the inset is adjusted to show that there is gold present in those particles. The scale bars represent 500 nm.

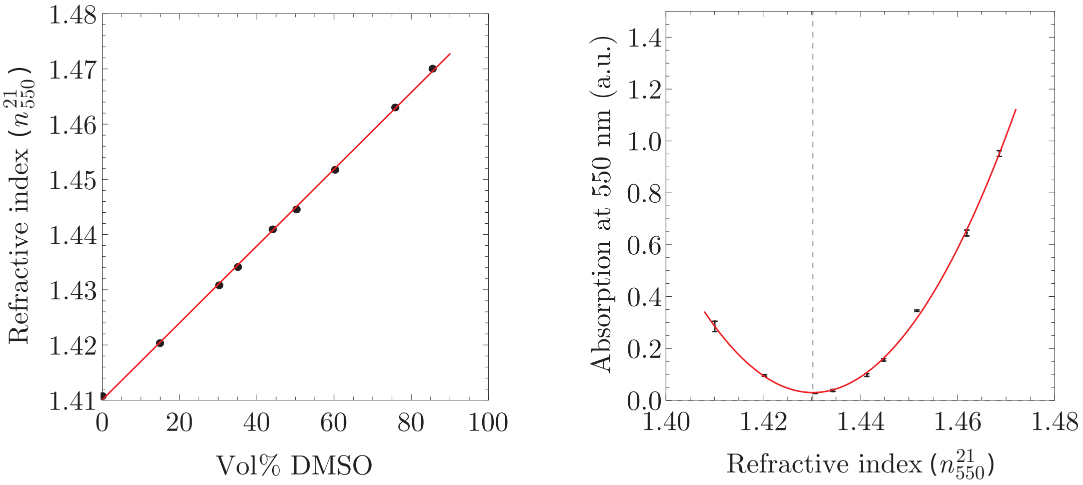


Supplementary Figure 7: In the left image, the refractive index at 550 nm and 21 °C ($n^{21}_{550}$) is plotted as a function of the vol% of DMSO in 1-pentanol. A linear equation was fitted to the data (red line). In the right image, the percentage of light transmitted at 550 nm is plotted as a function of the refractive index measured at 21 °C ($n^{21}_{550}$). The black dots correspond to measurements of dispersions of particles in DMSO/1-pentanol mixtures with various refractive indices. The error bars denote the standard deviation of the three absorption measurements for every refractive index. A quadratic equation was fitted to the data (red line) to determine the refractive index of maximum transmittance: $n^{21}_{550} = 1.4303$.

#### 2 Supplementary Tables

| Figure | Imaging Mode | Objective | PH (AU) | pixel sizes (nm) X / Y / Z | PDT (μs) | LAc | FAc | LAv | FAv |
|---|---|---|---|---|---|---|---|---|---|
| 2a | Confocal | 93× glyc | 0.7 | 35 / 35 / 50 | 2.0 | 2 | 4 | 1 | 1 |
| 3a | Confocal | 100× oil | 0.7 | 31.4 / 31.4 / 50 | 7.8 | 6 | 6 | 1 | 1 |
| 3b (l) | Reflection | 100× oil | 4.7 | 15 / 15 / - | 6.5 | 1 | 1 | 1 | 10 |
| 3b (r) | 2D STED | 100× oil | 1.0 | 20.5 / 20.5 / - | 4.3 | 6 | 4 | 1 | 1 |
| 3c (l) | Reflection | 93× glyc | 4.7 | 17 / - / 17 | 3.9 | 1 | 1 | 1 | 10 |
| 3c (r) | 3D STED | 93× glyc | 1.0 | 41 / - / 41 | 4.3 | 4 | 12 | 1 | 1 |
| 3d (l) | Reflection | 93× glyc | 4.7 | 20 / - / 20 | 6.5 | 1 | 1 | 1 | 10 |
| 3d (r) | 3D STED | 93× glyc | 1.0 | 41 / - / 41 | 4.3 | 4 | 12 | 1 | 1 |
| 4 (top)† | Reflection | 93× glyc | 1.0 | 20 / 20 / 80 | 3.8 | 1 | 1 | 1 | 1 |
| 4 (middle) | Confocal | 93× glyc | 1.0 | 20 / 20 / 98 | 0.3 | 1 | 4 | 2 | 1 |
| 4 (bottom) | 3D STED | 93× glyc | 0.6 | 20 / 20 / 98 | 0.3 | 1 | 4 | 2 | 1 |
| 5a | 2D STED | 100× oil | 1.0 | 11.3 / 11.3 / - | 2.2 | 2 | 1 | 1 | 8 |
| 5b | 2D STED | 100× oil | 1.0 | 11.3 / 11.3 / - | 2.2 | 4 | 1 | 1 | 8 |
| 5c | Reflection | 100× oil | 1.0 | 19.2 / 19.2 / - | 4.3 | 2 | 1 | 1 | 12 |
| 5e | 2D STED | 100× oil | 1.0 | 11.3 / 11.3 / - | 2.2 | 2 | 1 | 1 | 8 |
| 5f | 2D STED | 100× oil | 1.0 | 11.3 / 11.3 / - | 2.2 | 4 | 1 | 1 | 8 |
| 5g | Reflection | 100× oil | 1.0 | 19.1 / 19.1 / - | 4.3 | 2 | 1 | 1 | 12 |
| SI 6 | Reflection | 93× glyc | 1.0 | 21 / 21 / 97 | 1.28 | 1 | 1 | 1 | 25 |

Supplementary Table 1: Acquisition parameters of microscope images in Figs. 2-5 and Supporting Info Fig. 6. Key: 93× glyc = Leica HC PL APO 93×/1.30 GLYC motCORR STED WHITE, 100× oil = Leica HC PL APO 100×/1.40 OIL STED WHITE, PH = pinhole, AU = Airy units, PDT = pixel dwell time, L/FAc = Line/Frame Accumulation, L/FAv = Line/Frame Averaging.
† Stack average 10.

| Height (μm) | CC Position (%) |
|---|---|
| 0 | 44.5 |
| 20 | 40.5 |
| 40 | 38.5 |
| 60 | 33.4 |
| 80 | 20.0 |
| 100 | 11.1 |

Supplementary Table 2: Correction Collar (CC) positions chosen to equalize the axial 3D STED lobe intensities. These were set while recording the images in Fig. 4 of the main text.

### 3 Supplementary Notes

#### 3.1 Supplementary Note 1: Focused Ion Beam Scanning Electron Microscopy measurements

**FIB-SEM tomography**
In focused ion beam scanning electron microscope (FIB-SEM) tomography [2] a focused ion beam (FIB) scans a focused beam of Ga+ ions across the sample. The momentum transfer of the Ga+ ions sputters away atoms from the sample, allowing to mill cross sections with typical depths and widths of some tens of microns [3]. Subsequently, the cross sections can be directly imaged in-situ by the scanning electron microscope (SEM). The automated FIB-SEM tomography routine consists of alternately FIB milling away thin consecutive slices and SEM imaging of the resulting cross sections. This routine results in a series of images that can be reconstructed into a 3D representation of the analysed volume.

**Sample preparation**
A colloidal crystal with slightly larger particles (d = 531 nm, $< 2\%$ PDI) than used in the calibration sample was studied using FIB-SEM tomography. After growth, the colloidal crystal was infiltrated with the resin Lowicryl HM20. Infiltration of the crystal improves the finish of the milling process. Moreover, embedding the individual particles into a matrix prevents the particles to fall out of the crystal whilst slicing. The resin was composed of a 3:17 weight ratio mixture of Crosslinker D (Electron Microscopy Sciences) and Monomer E (Electron Microscopy Sciences). As initiator 0.5 w% dibenzoyl peroxide (Merck) was added. The crystal was infiltrated with resin which was subsequently cured at 65°C in an oven overnight.
The glass slide with the embedded crystal was attached to an aluminium SEM stub with carbon tape. A strip of carbon tape made contact between the crystal on top of the glass slide and the SEM stub to create an electrically conductive pathway. Subsequently, the ensemble was coated with a 5 nm thick layer of platinum, using a Cressington HQ280 sputter coater.

**Data acquisition**
A standard FIB-SEM tomography (Scios, AutoSliceAndViewG3 1.7.2, Thermo Fisher, former FEI) routine was carried out after protecting an area of interest with in-situ platinum deposition ($\sim$1 µm thickness) and preparing side trenches and an alignment marker. Milling conditions during the tomography run were 30 kV, 300 pA. The slice thickness was 75 nm. Images were recorded in BSE mode with 3.5 kV, 25 pA. The scan resolution was $3904 \times 4096$ which corresponded to a (horizontal) pixel size of 6.2 nm. The pixel dwell time was 10 µs. The series were recorded with the build-in auto-focus routine to ensure a high quality focus during the series.
The coordinates of the particles in the crystal were determined from the FIB-SEM tomogram as described in ref. [4].

#### 3.2 Supplementary Note 2: Refractive index matching to minimize scattering of the sample

To optimize the imaging conditions, the colloidal crystals should be infiltrated with a liquid with the same refractive index as the silica spheres. This refractive index matching minimizes scattering by the silica spheres as can be seen from the formula for the scattering of silica spheres in the Rayleigh-Gans-Debye approximation [5]:

$$I(K) \propto |m-1|^2 C_n R^6 P(K), \tag{1}$$

where $m = n_p/n_l$, $n_p$ and $n_l$ are the refractive indices of the particles and liquid, respectively, $C_n$ the particle number density, $R$ the particle radius and K the scattering vector:

$$K = \frac{4\pi n_l}{\lambda_0} \sin(\theta/2), \tag{2}$$

where $\lambda_0$ the wavelength of incident light in vacuum and $\theta$ the scattering angle.

We determined the refractive index of the silica particles by measuring the turbidity of a dispersion as function of the refractive index of the dispersion medium. Because this refractive index is temperature and wavelength dependent [6] this study was carried out under the same conditions as were used during confocal/STED measurements.

Therefore, all measurements were carried out at 21 °C, the estimated temperature in the confocal/STED setup, and transmittance was determined at 550 nm, the excitation wavelength used for confocal measurements.

The refractive index of the particles was determined by dispersing a fixed amount of particles in solutions with increasing refractive index. These solutions were prepared by mixing two solvents with different refractive indices in various ratios [7]. Next, the transmittance of these solutions was determined because it will reach a maximum when the refractive index of the particles and the solution match. The region of refractive indices that was studied, was chosen to be around $n_D = 1.45$ because for Stöber silica, $n_D$ values ranging from 1.43 to 1.462 are reported in literature [5, 8]. Two non-volatile solvents with refractive indices above and below 1.45 were selected for this: dimethyl sulfoxide or DMSO ($n_D^{25} = 1.4768$ [9]) and 1-pentanol ($n_D^{20} = 1.4103$ [10]).

**Materials**
Dimethylsulfoxide or DMSO (99.99%) and 1-pentanol (99%) were purchased from Sigma Aldrich and used as received.

**Methods**
First, the relationship between the vol% DMSO in 1-pentanol and the refractive index at 550 nm and 21 °C ($n_{550}^{21}$) was determined. To do so, a range of solutions going from 0 to ∼85 vol% DMSO in 1-pentanol to pure DMSO was prepared. With an Atago 3T refractometer, the $n_D^{21}$ and the corresponding Cauchy's equations of these solutions were determined. The Cauchy's equations were used to calculate the refractive index at 550 nm ($n_{550}^{21}$) [6].

For refractive index matching, 6.51 mL of a 1.92 vol% solution of particles in ethanol was transferred to a 20 mL glass vial and the ethanol was evaporated by placing this vial in a 50 °C preheated oven for 16 hours. Next, the particles were redispersed in 5 mL 1-pentanol by sonication to obtain a 2.5 vol% stock solution of particles in 1-pentanol. 0.5 mL of the stock solution of particles and various volumes of DMSO and 1-pentanol were transferred to separate vials to obtain a range of 0.25 vol% solutions of particles dispersed in different 1-pentanol/DMSO ratios (0 - 85 vol%). Absorption spectra of these solution were measured at 21 °C to determine the transmittance at 550 nm.

**Results**
In the left side of Supplementary Fig. 7 the refractive index at 550 nm and 21 °C ($n_{550}^{21}$) is plotted as a function of the vol% DMSO in 1-pentanol. A linear equation was used to fit this data. This linear relationship was used to calculate the refractive index of the particle suspensions that were used for the refractive index matching study. On the right side of the figure, the outcome of this study is presented. In this graph the transmittance at 550 nm and 21 °C of suspensions of particles with different refractive indices is plotted. By fitting a quadratic equation to this data, the refractive index of maximum transmittance was determined: $n_{550}^{21} = 1.4303$.
